# Inference of clonal selection in cancer populations using single-cell sequencing data

**DOI:** 10.1101/465211

**Authors:** Pavel Skums, Vyacheslau Tsivina, Alex Zelikovsky

## Abstract

Intra-tumor heterogeneity is one of the major factors influencing cancer progression and treatment outcome. However, evolutionary dynamics of cancer clone populations remain poorly understood. Quantification of clonal selection and inference of fitness landscapes of tumors is a key step to understanding evolutionary mechanisms driving cancer. These problems could be addressed using single cell sequencing, which provides an unprecedented insight into intra-tumor heterogeneity allowing to study and quantify selective advantages of individual clones. Here we present SCIFIL, a computational tool for inference of fitness landscapes of heterogeneous cancer clone populations from single cell sequencing data. SCIFIL allows to estimate maximum likelihood fitnesses of clone variants, measure their selective advantages and order of appearance by fitting an evolutionary model into the tumor phylogeny. We demonstrate the accuracy and utility of our approach on simulated and experimental data. SCIFIL can be used to provide new insight into the evolutionary dynamics of cancer. Its source code is available at https://github.com/compbel/SCIFIL

## 1 Introduction

Cancer is responsible for ~ 25% of deaths in the USA annually [19]. It is a disease driven by the uncontrolled growth of cancer cells having series of somatic mutations acquired during the tumor evolution. Cancer clones form complex heterogeneous populations, which include multiple clonal subpopulations constantly evolving to compete for resources, metastasize, escape immune system and therapy [11, 21, 14, 40]. Clonal heterogeneity plays key role in tumor progression [28, 26], and has important implications for diagnostics and therapy, since rare drug resistant variants could become dominant and lead to relapse in the patient [11, 8, 23]. Therefore cancer is now viewed as a dynamic evolutionary processes defined by complex interactions between clonal variants, which include both competition and cooperation [14, 40, 3].

Recent advances in sequencing technologies promise to have a profound effect on oncological research. Next-generation sequencing (NGS) already allowed for creation of large publicly available databases of cancer genomic data such as The Cancer Genome Atlas [37]. Study of genomic data for different tumors led to progress in understanding evolutionary mechanisms of cancer [40, 14, 21]. Most of cancer data have been obtained using bulk sequencing, which produces admixed populations of cells. However, the most promising recent technological breakthrough was the advent of *single cell sequencing* (scSeq), which allows to access cancer clone populations at the finest possible resolution. scSeq protocols combined with NGS allow to analyze genomes of individual cells, thus providing deeper insight into biological mechanisms of tumor progression.

The cornerstone of such analysis is an estimation of parameters defining the evolution of heterogeneous clonal populations. Currently there is no scientific consensus about the rules guiding the evolution of cancer cells [6, 35, 39, 30], with multiple competing theories being advanced by different researchers. The open questions include the rules of evolution (neutral, linear, branching or punctuated), ways of interaction between members of heterogeneous populations (competition or cooperation) and the role of epistasis (non-linear interaction of different SNVs or genes). These questions could be addressed by estimation of evolutionary parameters for cancer lineages from NGS data [39, 35].

One of the most important evolutionary parameters is the collection of replicative fitnesses of individual genomic variants, commonly termed *fitness landscape* in evolutionary biology [13]. Several computational methods have been proposed for *in vitro* estimation of fitness landscapes [33, 25, 16, 12]. However, *in vitro* studies are cost- and labor-intensive, consider organisms removed from their natural environments and does not allow to capture all population genetic diversity [34]. One of the possible ways to infer fitness landscape *in vivo* is to analyze follow-up samples taken from a patient at multiple time points and compute fitnesses directly by measuring changes of frequencies of genomic variants over time. However, follow-up samples are very scarce, and overwhelming majority of data represent individual samples.

Quantification of clonal selection from individual samples is computationally challenging, but extremely important for understanding mechanisms of cancer progression [39, 35]. In particular, recent findings on structures of fitness landscapes of cancer from bulk sequencing data [38] initiated a lively scientific discussion published in several papers [35, 30, 39]. It can be anticipated that single cell sequencing data will be able to shed light into this important problem. It is known that relative abundances of genomic variants alone are not indicative of variant fitnesses [34]. Existing methods for inference of fitnesses from single samples utilize more sophisticated approaches, but have various limitations including reliance on the assumption that the population is in equilibrium state, or disregard of population heterogeneity and variability of fitness landscapes, or customization to bulk sequencing data [34, 7, 39].

### Contributions

We propose a computational method SCIFIL (Single Cell Inference of FItness Landscape) for *in vivo* inference of clonal selection and estimate of fitness landscapes of heterogeneous cancer clone populations from single cell sequencing data. SCIFIL estimates fitnesses of haplotypes rather than alleles, and does not assume allele independence which allows to take into account the effects of epistasis. Instead of assuming that sampled populations are in the equilibrium state, our method estimates fitnesses of individual clone types using maximum likelihood approach. We demonstrate that the proposed method allows for accurate inference of fitness landscapes and quantification of clonal selection. We conclude by applying SCIFIL to real tumor data.

## 2 Methods

We propose a maximum likelihood approach, which estimates fitnesses of individual haplotypes by fitting into the tumor phylogeny an evolutionary model with the parameters explaining the observed data with the highest probability. We first establish the ordinary differential equations (ODE) model for the tumor evolutionary dynamics, and define the likelihood of the observed data given the model parameters. We conclude with finding fitnesses maximizing the likelihood by reducing the problem to finding the most likely mutation order and applying branch-and-bound search to solve that problem.

Traditionally, evolutionary histories are represented using binary phylogenetic trees. Following [18], we use an alternative representation of an evolutionary history of a tumor using a *mutation tree*. The internal nodes of a mutation tree represent mutations, leafs represent single cells, internal nodes are connected according to their order of appearance during the tumor evolution and the mutation profile of each cell equals the set of mutations on its path to the root (Fig. 1). In addition we accumulate all leafs attached to the same internal node into a single leaf with an abundance representing a particular haplotype. For simplicity of further calculations we assume that there is a leaf attached to every internal node, with some leafs having an abundance 0 (or rather some small number *δ* << 1). Unlike [18], generally we do not need to employ the infinite site assumption (or that the mutation tree represent a *perfect phylogeny*), i.e. repeats of mutations are allowed provided that mutation profiles of all haplotypes in a tree are unique. It agrees with recent findings [22]. A mutation tree can be constructed using currently available tools, such as SCITE (without [18] or with [22] infinite site assumption) or SiFit [41].

**Figure 1:**
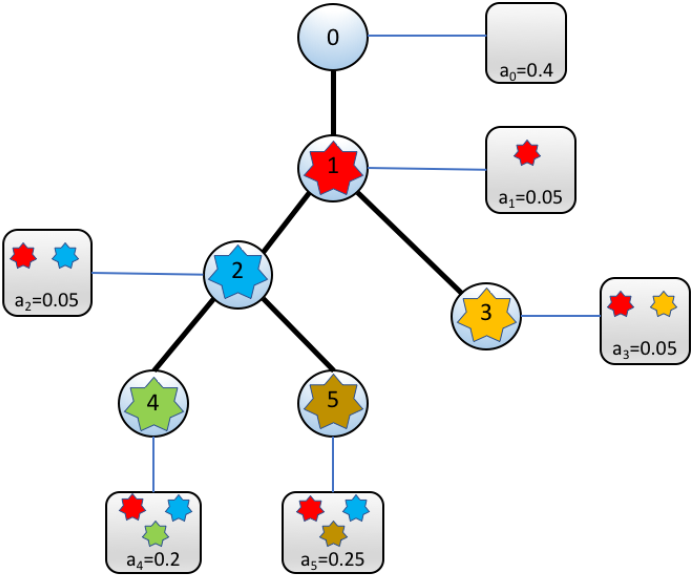
Mutation tree

Formally, we consider the following algorithmic problem. Given are:

- mutation tree *T* with *n* + 1 leafs corresponding to haplotypes. We assume that internal nodes of *T* are labeled 0, 1, …, *n* and the *ith* haplotype is attached to the node *i*. The root of *T* correspond to the mutation 0, which represent absence of somatic mutations or healthy tissue.
- observed relative abundances 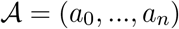 of haplotypes.
- Mean cancer cells mutation rate *θ*. It is well-studied evolutionary parameter with estimations provided by various prior studies [36, 15].

The goal is to find haplotype fitnesses 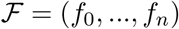 that maximize the likelihood

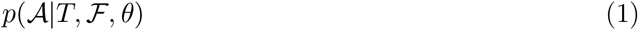

The current section is organized as follows. First we introduce our evolutionary model of choice and the general definition of the probability (1). Next, we describe how the likelihood is modified to transform the maximum likelihood problem (1) into a discrete optimization problem. Finally, we describe the proposed method of estimation of fitnesses 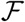 maximizing (1).

### Evolutionary model

We consider tumor evolution as a branching process described by the mutation tree *T*. Let *V*(*T*), *V_I_*(*T*) and *E*(*T*) be node set, internal node set and an arc set of *T*, respectively. Let also *p_i_* denote the parent of a node *i* ∈ *V_I_*(*T*). We assume that nodes *V_I_*(*T*) represent mutation events, with *j*th event occurring at rate *θ_j_*. The mutation event corresponding to a node *i* happens at time *t_i_*; at the event the haplotype *p_i_*, gives birth to a haplotype *i*. The dynamics of the cancer clone population is described by the piecewise continuous function *x* = (*x*_0_, …, *x_n_*), where *x_i_* = *x_i_*(*t*) is the relative abundance of the *i*th haplotype. The discontinuity points of *x* correspond to mutation events. Let *r, i, j* be 3 consecutive mutation events with times *t_r_* < *t_i_* < *t_j_*. Between mutation events *i* and *j* at interval [*t_i_, t_j_*] haplotype frequencies 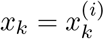 follows the system of ODEs [32]:

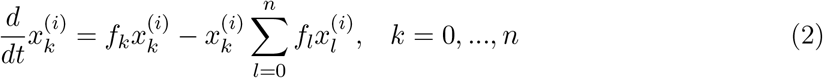

with initial conditions

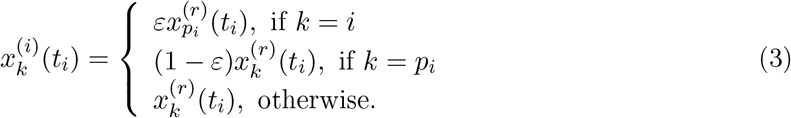

Here subtraction of the term 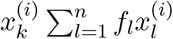 in (2) ensures that the total relative abundance of haplotypes sum up to 1. Initial conditions (3) link haplotype abundances before and after the mutation event *i* and indicate that at time *t_i_* the haplotype *i* is generated by the haplotype *p_i_*. The parameter *ε* is a small number (by default *ε* = 10^−5^). At time 0, the root haplotype (healty tissue) gives birth to the first mutation, with the corresponding haplotypes having relative abundances 1 − *ε* and *ε*.

### Likelihood definition

In addition to *n* mutation events, we consider the (*n* + 1)th event representing haplotype sampling. Suppose that times of mutation events 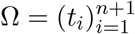 and mutation rates between events 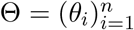 are given. Let *σ* = (*σ*_1_, …, *σ*_*n*+1_) be the permutation of events in order of their appearance, i.e. 0 = *t*_*σ*_1__ < *t*_*σ*_2__ < … < *t_σ_n__* < *t*_*σ*_*n*+1__). The probability of observing abundances 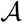 given *T*,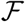, Ω, Θ and *θ* is defined as the product of probabilities of mutation events and probabilities of observed haplotype abundances.

Consider the mutation event in the vertex *σ_j_*. The event occurs if 2 conditions are met:

a. no mutation events have been observed over the time interval (*t*_*σ*_*j*−1__, *t_σ_j__*);
b. at time *t_σ_j__* the mutation happened in the haplotype *p_σ_j__* rather than in other haplotypes which exist at that time.

Appearance of mutation is a classical rare event, and therefore we assume that the time intervals between consecutive mutation events *i* and *j* follow a Poisson distribution with the mean 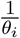. Mutation rates are distributed normally with the mean *θ* and the standard deviation *ν*. Assuming that mutations are random, the probability of (b) is equal to the frequency *x_p_σ_j___*(*t_σ_j__*) of the haplotype *p_σ_j__* at time *t_σ_j__* according to the system (2). Finally, we assume that the probability of seeing observed frequencies given model-based frequencies at the sampling time follows a multinomial distribution 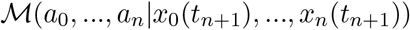. After putting all probabilities together, we have

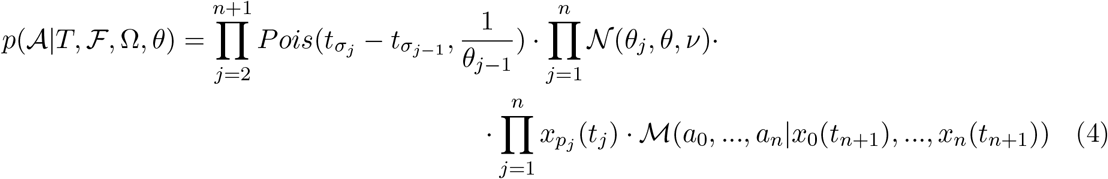

Our goal is to find best fitting fitnesses 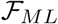, rates *Θ_ML_* and times Ω_*ML*_ by solving the following maximum likelihood problem:

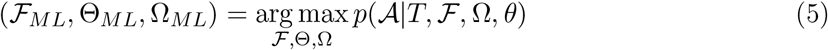

The probabilities 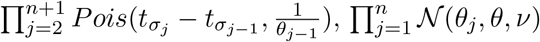, 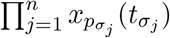 and 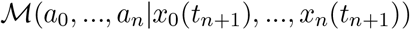 are further referred as *time likelihood, rate likelihood, mutation likelihood* and *abundance likelihood*, respectively. For the tree shown on Figures 1 and 2 it is equally feasible that the mutation 2 appeared before the mutation 3 or vise versa. However, haplotype 2 later produces mutations 4 and 5, and therefore the mutation likelihood suggests that at that mutation events it had high abundance. This situation is probable if either 2nd mutation appeared earlier or it appeared later but has a high fitness. Time, rate and abundance likelihoods allow to choose between these two alternatives.

**Figure 2:**
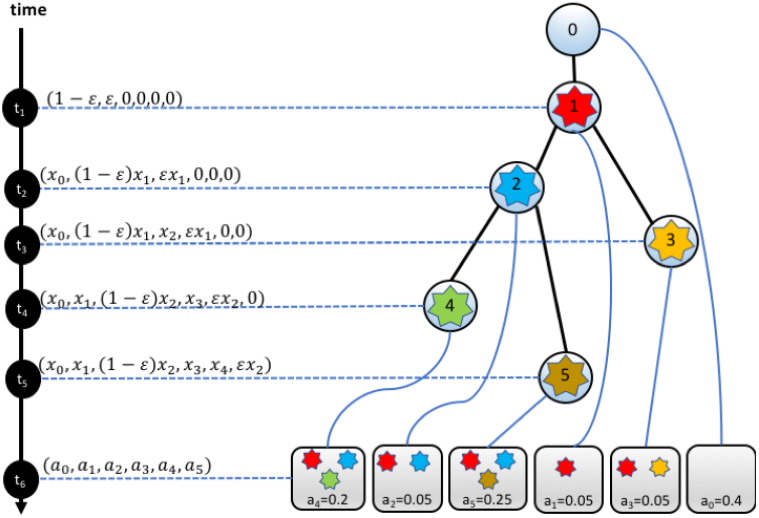
Illustration of the evolutionary model. Each node of the tree represent a mutation event whose times are marked on the time axis. Leafs represent the sampling event. Next to each time point the distribution of haplotype abundances after the corresponding event is shown.

### Reduction to discrete optimization

The standard way to solve the maximum likelihood problem (5) is to optimize 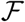, Θ and Ω jointly using Markov Chain Monte Carlo (MCMC) sampling. However, our experiments have shown that the function (1) has too many local optimums which makes MCMC search over the continuous space of possible solutions inefficient. Therefore we suggest an alternative heuristic approach, which transforms the problem (5) into a discrete optimization problem akin to a scheduling problem. This problem is then solved using a specifically designed combinatorial heuristic search.

Firstly, we assume that all fitnesses are relative with respect to a fitness of a haplotype 0 which is set to be *f*_0_ = 1. By default, this haplotype corresponds to the normal tissue. For the problem of inference of clonal selection such assumption does not restrict the predictive power. Next, we observe that any assignment of event times Ω defines the order of appearance *μ_i_* for each node *i* ∈ *V*(*T*) (e.g. on Figure 2 *μ_i_* = *i* for *i* = 1, …, 5). This order agrees with the natural vertex order induced by *T*, i.e. *μ_i_* < *μ_j_* whenever *i* is an ancestor of *j*. In turned out that conversely any order *μ* defines times Ω^*μ*^, rates Θ^*μ*^ and fitnesses 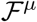 which maximize the partial likelihood

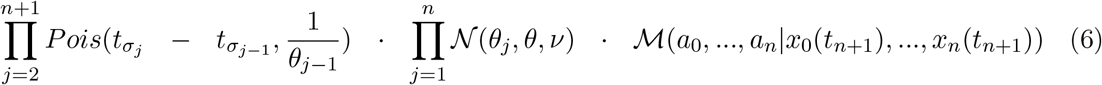

More precisely, the following proposition holds.

#### Proposition 1

*For a given order vector μ, times* Ω*^μ^, rates* Θ*^μ^ and fitnesses* 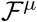 *maximizing (6) can be estimated as follows:*

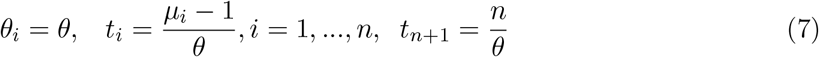

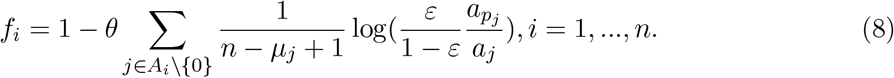

*Here A_i_ is the set of ancestors of a node i (including itself)*.

#### Proof

Poisson and Gaussian probabilities achieve maximums at their means, i.e. the rate and time likelihoods are maximal, when for consecutive events *i, j* we have 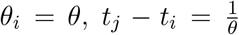. This yields the solution (7). The multinomial probability 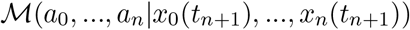 is maximal when *x_i_*(*t*_*n*+1_) = *a_i_* for all 0 = 1, …, *n*. This can be rewritten as

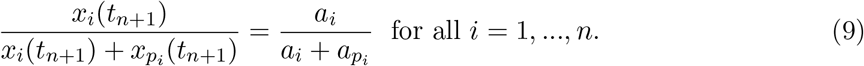

Our goal is to find fitnesses 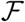 such that (9) holds. We find an approximate solution to this problem by disregarding the discontinuity of the abundances 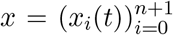. We use the observation that the system (2) is invariant with respect to the transition to relative abundances of any pair of haplotypes. Namely, for each haplotype pair *i, j* = 0, …, *n* dynamics of their relative abundances with respect to each other 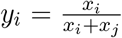 and 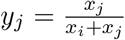 is described by the system of ODEs of the same form as (2):

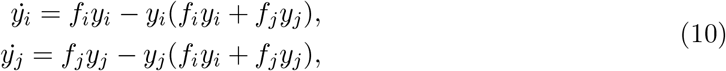

On the interval [*t_i_, t*_*n*+1_] relative abundance 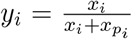 satisfy the system (10) with the initial condition *y_i_*(*t_i_*) = *ε*. After shifting time interval to [0, *t*_*n*+1_ − *t_i_*] this system can be linearized and solved in closed form, producing a solution

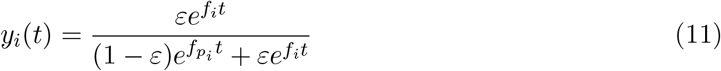

After putting the expressions (11) into the equations (9) with *t* = *t*_*n*+1_−*t_i_* we get the following system of equations to find fitnesses 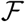:

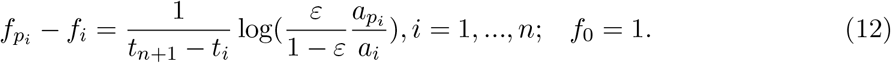

Solving it with *t_i_* described by (7) yields the solution (8).

Taking into account Proposition 1, we replace the maximum likelihood problem (5) with the following discrete problem: find the ordering *μ* maximizing the mutation log-likelihood

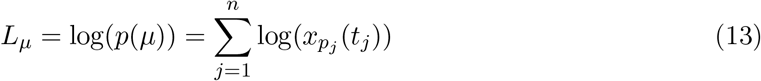

with times Ω^*μ*^ and fitnesses 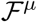 described by (7),(8) subject to the constraint that *μ* agrees with with the ancestral-descendant order of *T*.

### Finding optimal ordering

The problem (13) could be considered as a variant of scheduling problem with precedent constraints and with non-linear cumulative cost function [9]. Here mutations play roles of jobs, ordering of mutations corresponds to scheduling of jobs on a single processor, mutation tree represent job precedence constraints, and the objective (13) indicates that the cost of processing of a job depends on the previously processed jobs. Such problems are usually NP-hard [9]. We solve it by a heuristic approach combined with the search in the space of feasible sub-orderings of nodes of the mutation tree *T*. The general scheme of our method is described by Algorithm 1. The algorithm starts with the initial tree *T*′ = *T* and iteratively transforms it into a total order as follows. We call two simple paths of *T*′ *sibling paths*, if they share the starting vertex. We traverse the nodes of *T*′ in a bottom-up direction and merge sibling paths into one path representing optimal sub-order of their nodes with respect to the objective (13). The algorithm stops when all nodes form a single path.

Merging of sibling paths *P*_1_ and *P*_2_ is performed by Algorithm 2. We note that feasible orders of paths’ nodes bijectively correspond to *k*-subsets of a *k* + *l*-set [*k* + *l*]: for a given *k*-subset *X*, a feasible order *μ_X_* can be obtained by placing nodes from *P*_1_ \ {*u*} (resp., *P*_2_ \ {*u*}) at positions from *X* (resp., [*k* + *l*] \ *X*) in order of their appearance in *P*_1_ (resp., *P*_2_); inverse is also true. Algorithm 2 recursively generates *k*-subsets via branching and prune branches, if the corresponding orders are likely to be sub-optimal.

The *k*-subsets are generated recursively in a standard way [?] using the property that every *k*-subset *X* of [*k* + *l*] is either *k*-subset of the set [2: *k* + *l*] or has the form *X* = {1} ∪ *Y*, where *Y* is a *k* − 1-subset of [2: *k* + *l*]. Suppose that at a given iteration a partial *k*′-subset *Y, k*′ ≤ *k*, and the corresponding pre-order *μ_Y_* has been constructed. For all nodes *v* covered by *μ_Y_* we calculate their appearance times *t_v_* and fitnesses *f_v_* using (7),(8), and abundance distributions *x^v^* = (*x*_0_(*t_v_*), …, *x_n_*(*t_v_*)) from the system (2)-(3) (in fact, it is not necessary to recalculate all values since some of them has been already calculated at previous iterations). Next, we heuristically extend *μ_Y_* to a total order as described below. If the likelihood of the constructed solution is below the current optimum, then the recursion tree branch of the partial solution *Y* is pruned. Otherwise, the current optimum is updated and the recursion continues.

#### Algorithm 1 Algorithm for node ordering

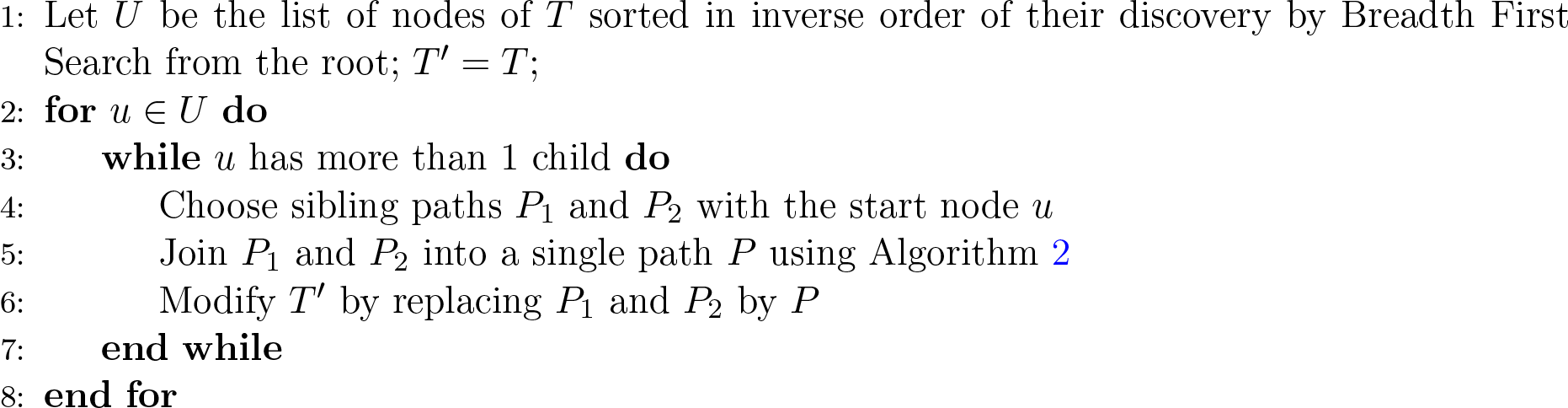

#### Algorithm 2 Algorithm for path joining

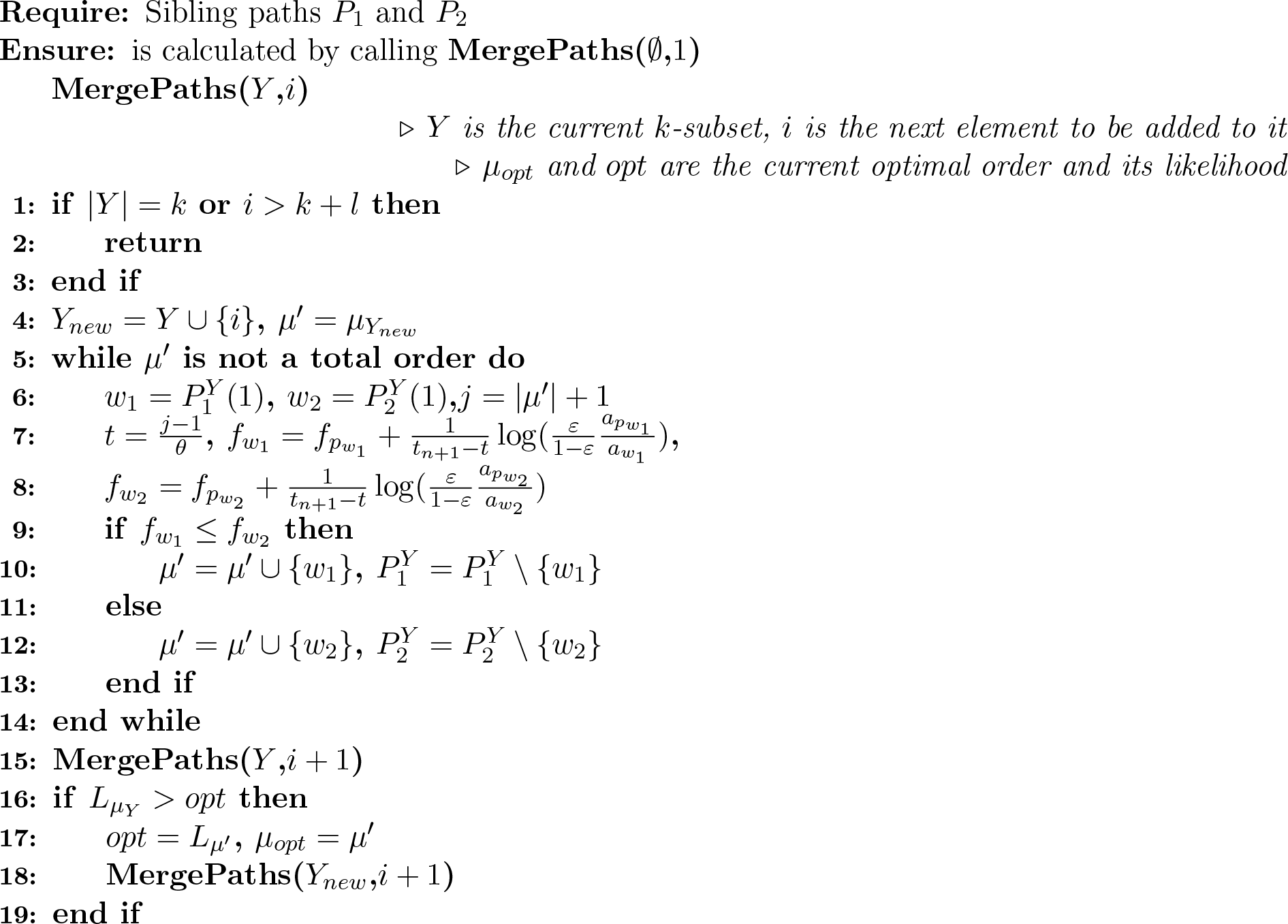

Finally, we describe how an order *μ_Y_* is extended (lines 5-14 of Algorithm 2). We consider the subpaths 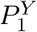 and 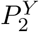 formed by the nodes of *P*^1^ and *P*^2^ that are not covered by *μ_Y_*. For the first nodes of these subpaths, we calculate their provisional fitnesses under the assumption that each node is added to *μ_Y_* as the next element. The node with the smaller provisional fitness is added to *μ_Y_*. This procedure is repeated until *μ_Y_* covers all nodes. The logic behind this approach is based on the observation that according to (2) the frequency of a haplotype grows while its fitness is larger than the average fitness of the population, and declines otherwise. For a given iteration, adding haplotype with a smaller fitness slows down the average fitness growth. As a result, for preceding haplotypes probabilities of appearances of their children in the future may become higher.

## 3 Results

### 3.1 Simulated data

We simulated 100 test examples with *m* = 30, 60, 90, 120, 150 mutations, which correspond to numbers of mutations for real single cell sequencing data analyzed in previous studies [18, 21, 24]. For each test example, clonal evolution was simulated as follows. (a) Mutations 1, …, *m* are generated randomly. For the time interval between mutation events *i* and *i* + 1 the current mutation rate *θ_i_* is sampled from the normal distribution with the mean *θ* = 0.01 and standard deviation *σ* ∈ {0.1 · *θ*, 0.5 · *θ*, 0.9 · *θ*}. At each moment of time of that interval a mutation event happens with the probability *θ_i_*; at the event a random haplotype *p* selected with the probability equal to its current relative abundance gives birth to a new haplotype *j* with the random fitness *f_j_* by acquiring a random mutation *i* + 1. (b) When there is no mutation event, abundances of existing haplotypes are updated according to the model (2). After the end of the simulation, final abundances were randomly perturbed by 10% to incorporate the possible noise in their estimation. The simulated mutation tree and haplotype abundances were used as an input for SCIFIL. The tool was run with the timeout of 1hr.

We quantified the performance of SCIFIL using two performance measures:

1. Mean relative accuracy 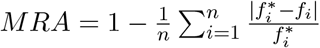, where 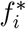 and *f_i_* are true and inferred fitnesses, respectively.
2. Spearman correlation *SC* between vectors of true and inferred fitnesses.

*MRA* and *SC* highlight different aspects of the problem. MRA measure the accuracy of fitness value estimation, while in our case SC measures how well we are able to qualitatively detect selective advantage of particular clones over other clones. In particular, when the population contains several clones with close fitnesses, even small error in fitness estimation may cause low SC. Correct fitness ranking can be used in evolutionary studies even when actual fitness values are missing or inaccurate [5].

The results of SCIFIL evaluation on simulated data are shown on Figs. 3-4. The algorithm demonstrated high accuracy as measured by both parameters. The number of mutations (Fig. 3) does not have a great impact on the Spearman correlation, which averages 97.35% (standard deviation 1.2%) over all analyzed test cases. MRA decreases when the number of mutations grows, but remains above 88% for all datasets. Increase in variation of mutation rate (Fig. 4) does not significantly affect SC, and results in slight decrease of average MRA. Relative robustness of SCIFIL to the variation of mutation rates (which also introduce variation in mutation times) indirectly suggests, that the proposed algorithm is able to well approximate the original maximum likelihood problem (4).

**Figure 3:**
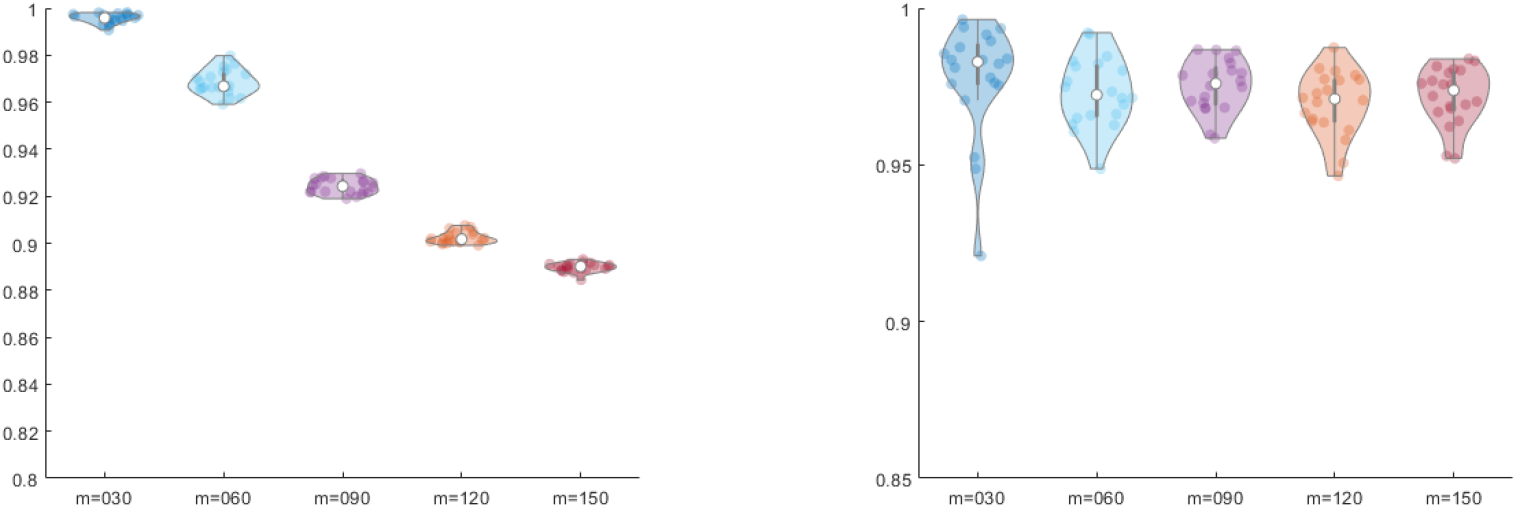
Performance of SCIFIL on simulated data with *m* = 30, 60, 90,120,150 mutations and fixed standard deviation of mutation rate. Left: violin plots of the mean relative accuracy of fitness estimation. Right: violin plots of Spearman correlation between true and inferred fitness vectors

**Figure 4:**
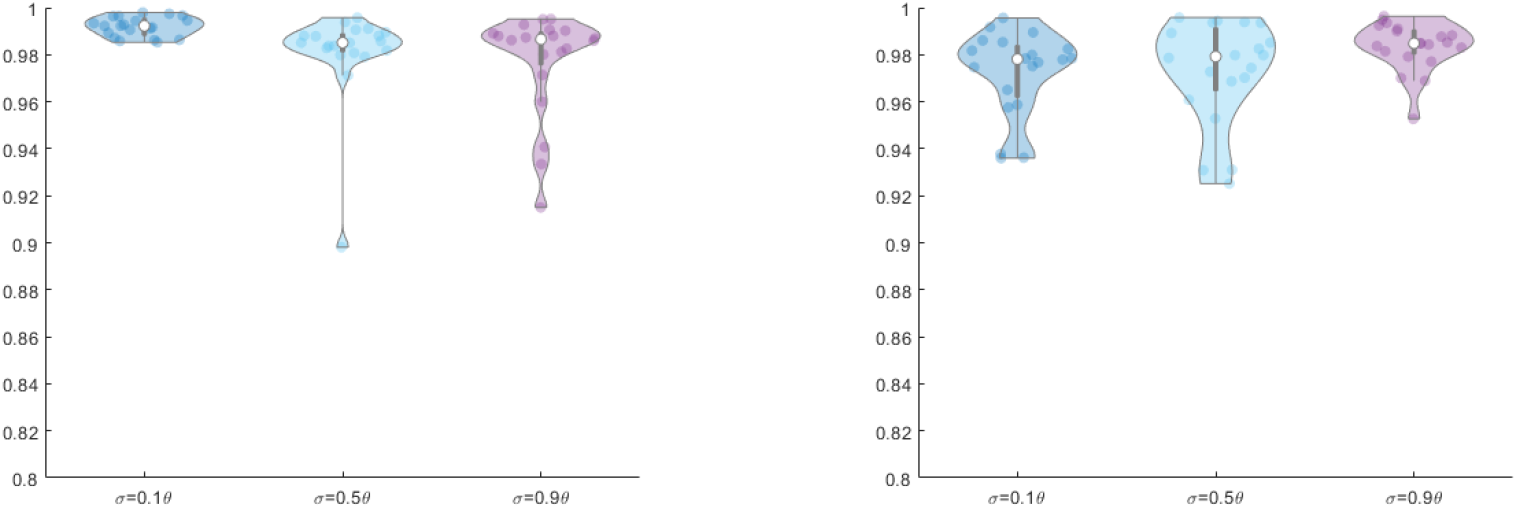
Performance of SCIFIL on simulated data with *m* = 50 mutations and different standard deviations of mutations rates. Left: violin plots of the mean relative accuracy of fitness estimation. Right: violin plots of Spearman correlation between true and inferred fitness vectors

We compared our approach with the previously published tool QuasiFit [34]. Although originally designed for viruses, QuasiFit is based on quasispecies model, which is applicable to both intra-host viral populations and cancer clone populations [10] and is essentially a fully continuous version of the model used by SCIFIL. Both QuasiFit and SCIFIL reconstruct replicative fitnesses of individual haplotypes (rather than alleles). In addition to genomic data, both algorithms utilize other information: SCIFIL uses a mutation tree, while QuasiFit assumes that the population is in equilibrium state of the quasispecies model. Furthermore, SCIFIL is a discrete optimization approach, while QuasiFit implements Markov Chain Monte Carlo sampling. QuasiFit was run with the per-cell mutation rate *μ* = *εθ* (which is a fully continuous analogue of the parameters used by SCIFIL) and inferred fitnesses were considered after a burn-in of 100000 iterations. As QuasiFit uses a different fitness vector normalization, following [34] we used only the parameter *SC* for the comparison. The results are shown on Fig. 5 (left). SCIFIL outperforms QuasiFit indicating that in certain settings the proposed model could be more accurate for the inference of clonal selection than the equilibrium state assumption. For the sake of fairness, it should be noted that unlike QuasiFit SCIFIL has access to information about partial haplotypes order encoded by the mutation tree, which facilitates correct estimation of fitness correlation.

**Figure 5:**
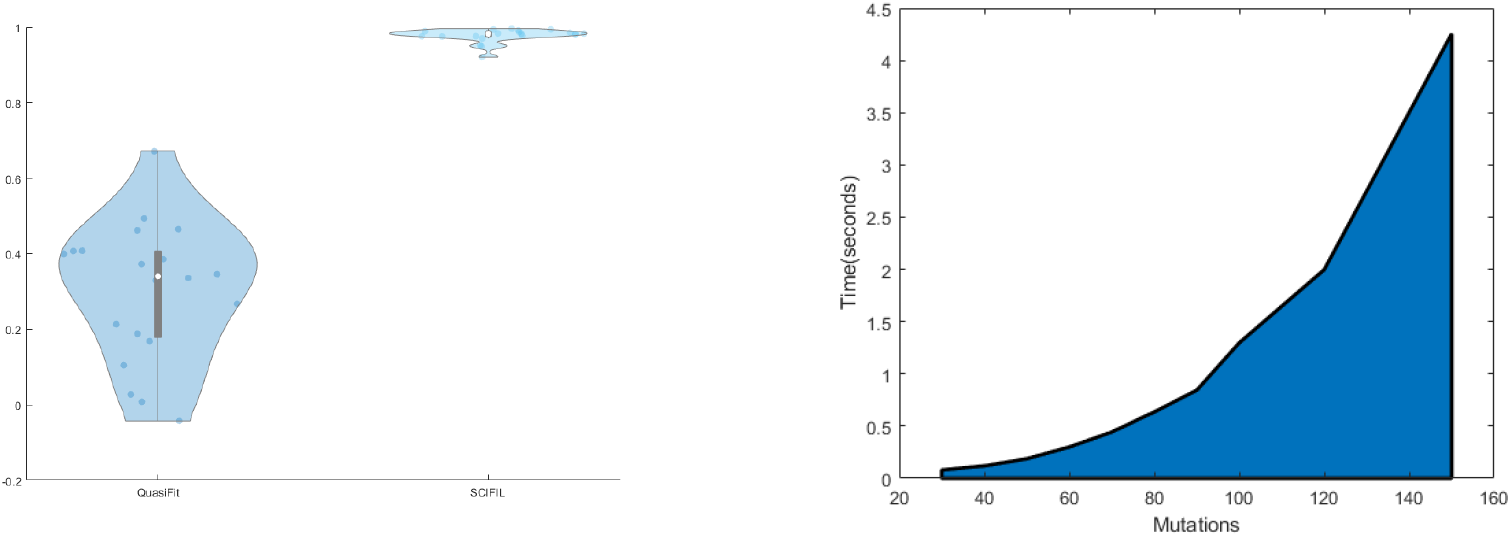
Left: Spearman correlation between true and inferred fitness vectors for QuasiFit and SCIFIL. Right: running time of SCIFIL

Computational experiments suggest that the algorithm’s running time scales quadratically with the number of mutations (Fig. 5 (right)). It allows SCIFIL to finish in a few seconds for all analyzed data sets when run on a simple desktop computer.

### 3.2 Experimental data

We used SCIFIL to infer fitness landscapes for several recently published experimental cancer datasets. Due to the page restrictions, here we present the results for 2 datasets which are particularly illustrative. The first dataset is single-cell sequencing data from a JAK2-negative myeloproliferative neoplasm (essential thrombocythemia) [17, 18], the second one represents metastatic colon cancer [24]. The latter dataset includes SNVs sampled from the main tumor and two metastases. We confined our analysis only to the primary tumor, since it is biologically meaningful to compare fitnesses of clones sampled from the same environment. For both datasets, their mutation trees were reconstructed using SCITE [18], and fitnesses and mutation appearance times were inferred by SCIFIL with the cell-wise mutation rate 10^−6^. It is important to note that varying SCIFIL parameters may change absolute values of inferred fitnesses, but preserve relations between them. The relations are the most informative factors for evolutionary analysis.

We visualized inferred fitness landscapes as follows. First, we calculated a distance between each pair of haplotypes defined as the sum of their hamming distance and the absolute difference of their orders of appearance. Obtained distances were used to map haplotypes to the plain ℝ^2^ using multidimensional scaling. Fitness values of the points corresponding to haplotypes were interpolated using biharmonic splines, and the resulting surface was visualized as a contour plot with the tree placed on top of the plot. The obtained plots are shown on Fig. 6, where colors represent fitness values, and distance from each tree node to the root reflects its appearance time.

**Figure 6:**
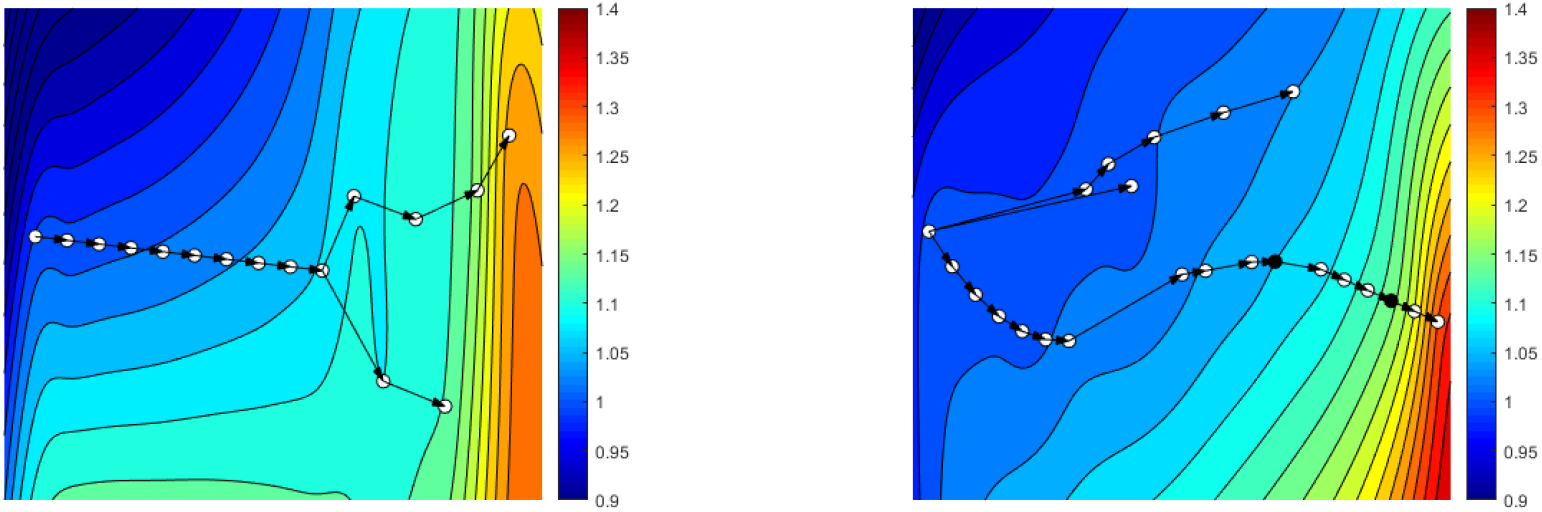
Fitness landscape and mutation tree for JAK2-negative myeloproliferative neoplasm [18, 17] (left) and colorectal cancer (right) [24] inferred by SCIFIL. Colors represent fitness values, and distance from each tree node to the root is approximately proportional to its time of appearance.

For myeloproliferative neoplasm (Fig. 6, left) we observe linear accumulation of mutations with slight selective advantages at the beginning of tumor evolution, followed by the subclone expansion of two lineages with significantly faster fitness growth. The rate of fitness growth after the branching event is ~ 3 times higher than before it. Thus, answering the question posed in [18], we may predict that recent subclones will replace ancestral clones. However, based on the available information it is hard to decide whether one of subclone lineages will out-compete another one, or they will continue to coexist.

Evolution of the colon tumor (Fig. 6, right) follows different scenario, with 3 independent lineages co-existing at the beginning without a clear selective advantage enjoyed by any of them. This stage is followed by the fast expansion of one of the lineages, which climbs a fitness peak and acquires selective advantage over other lineages. Exactly at this stage the advantageous lineage seeded the metastatic tumor at two seeding events (highlighted in black on Fig. 6).

Experimental data also allow to emphasize how SCIFIL estimations extends predictions implied by underlying evolutionary model. Although the model suggests positive selection with fitness growth along each path of the mutation tree as the most probable scenario, it does not imply any restrictions on the comparative fitnesses of different lineages. In particular, fitness advantages of haplotypes are not defined only by their distances from the root, as emphasized by the fitness landscape of the colon tumor, where, for instance, the node highlighted in purple has higher fitness than the node highlighted in red. The reason is that haplotype abundances contribute to the estimation of fitness values as much as the evolutionary model and the topology of mutation tree.

## 4 Discussion

Intra-tumor heterogeneity is one of the major factors influencing cancer progression and treatment outcome. Cancer clones form complex populations of genomic variants constantly evolving to compete for resources, proliferate, metastasize and escape immune system and therapy. Quantification of clonal selection and inference of fitness landscapes for individual tumors may provide valuable information for understanding mechanisms of disease progression and for design of personalized cancer treatment. Single cell sequencing provides an unprecedented insight into intra-tumor heterogeneity promising to allow to study fitness landscapes at finest possible resolution and quantify selective advantages on the level of individual clones.

In this paper, we presented SCIFIL, a likelihood-based method for inference of fitnesses of clonal variants. Unlike other available methods for related problems, SCIFIL takes full advantage of the information about structure and evolutionary history of clonal population provided by single cell sequencing. It uses individual cells as evolutionary units, in contrast to the tools based on bulk sequencing which perform their analysis on the level of subpopulations or lineages. Furthermore, SCIFIL can also handle bulk sequencing data as long as haplotypes are reconstructed and mutation tree is constructed using available tools.

The proposed approach differs from previous approaches by employing more general and realistic evolutionary model rather than assuming that the population achieved the equilibrium state. We have demonstrated that our approach allows for accurate inference of fitness landscapes and can be used for analysis of evolutionary history and clonal selection for real tumors. We envision that SCIFIL can be also used to infer epistasic interactions and to identify combinations of mutations and/or mutation patterns driving the tumor growth. In addition, it can be applied to other highly mutable heterogeneous populations, such as viral quasispecies or bacterial communities.

The proposed approach has limitations which should be taken into account and/or addressed in the future work. Fitness is not defined by the genetic composition alone, and depends on the environment. Thus SCIFIL quantitative predictions are more reliable when the analyzed clones are sampled from the same tumor. Fitness inference relies on the observed clone abundances, and therefore significant inaccuracies in abundance estimation may affect accuracy of fitness reconstruction. For single cell data it is particularly important owing to its susceptibility to allelic dropouts and PCR bias. However, this problem can be addressed by using a combination of bulk and single cell sequencing data. There exist plethora of tools which can estimate clone abundances from composite bulk and single cell sequencing data (see, e.g. [1, 29]). In addition, such composite data can be employed to increase an accuracy of mutation trees reconstruction [27]. We expect SCIFIL reliability to increase when it will be combined with these tools.

Another set of limitations arise from the selected evolutionary model. The model (2) was selected due to its generality [10] and suitability for fitness landscape inference [31]. However, it has certain underlying assumptions: the mutation rates are supposed to be normally distributed, while the dynamical system (2) implies positive selection with the gradual growth of average population fitness. It should be noted that in many cases such assumptions are sufficiently realistic or general, and has been used in several studies to obtain valuable insights into the dynamics of tumor evolution [4, 20]. In particular, other studies demonstrated that even a normal mutation rate is sufficient to produce significant intra-tumor heterogeneity and emphasized the relative importance of selection over both the size of the cell population and the mutation rate [2]. Although equations (12) suggest that in most cases fitness growths along each path of the mutation tree, the model does not imply any restrictions on the comparative fitnesses of different lineages. Furthermore, observed relative abundances of haplotypes are independent of the model, and their contribution to the estimated fitness values is paramount. Nevertheless, we expect that our approach can be extended by incorporating other models capturing different evolutionary scenarios, such as gradual mutation rate growth over the course of tumor evolution. It should be noted, though, that currently there is no universal evolutionary model for tumor progression. Alternative models will inevitably introduce other limitations and can be less practical for fitness estimation.

On algorithmic side, the optimization problem behind our approach can be viewed as the type of scheduling problem with precedent constraints and with non-linear objective [9]. Such problems are generally NP-hard, although the complexity of our problem is unknown. It is known that for certain simple objectives and well-structured precedence constraints (e.g. defined by series-parallel graphs) the corresponding scheduling problems are polynomially solvable [9]. For our problem precedence constraints have the form of a tree. It gives a certain hope of existence of exact polynomial or a good approximation algorithm, although the complex objective function may keep our problem NP-hard. This question requires additional study.

